# Extreme disparity in the appendicular skeleton of domestic dogs (*Canis familiaris*)

**DOI:** 10.64898/2026.03.22.713490

**Authors:** Lucy E Roberts, Olivia Binfield, James P Charles, Eithne J Comerford, Karl T Bates, Anjali Goswami

## Abstract

Domestic dogs (*Canis familiaris*) display more morphological variation than any other mammal. Cranial morphology has been extensively studied, as have the relationships with function, development, genetics, veterinary medicine, and breed welfare. Postcrania remain comparatively understudied, despite well-documented breed-specific predispositions to musculoskeletal disease. Here, we apply three-dimensional landmark-free morphometrics to quantify the shape of 743 elements from 213 dogs, including the scapula, humerus, radius, ulna, pelvic girdle, femur, tibia, and fibula. We assess integration among limb elements and investigate drivers of shape variation within and between breeds. Across most breeds, limb bone shape is strikingly similar. Dachshunds, however, exhibit distinct morphology across all elements and one to two orders of magnitude greater variation than any other breed. Despite this disparity, integration remains high between all element pairs. Remarkably, we find no significant relationship between bone shape and body mass, age, or pathology, but comparison with historic specimens reveals marked changes in dachshund long bone shape over the past ∼150 years. These extreme differences are not shared by other sampled chondrodysplastic breeds, underscoring the need to understand morphological diversity beyond simple categorisation. These findings provide a quantitative framework for linking postcranial morphology with function, disease risk, and evidence-based improvements to canine welfare.

## Introduction

The domestic dog (*Canis familiaris*) is often considered to be the most morphologically diverse species of mammal on Earth today. *C. familiaris* likely arose from a now-extinct population of grey wolves 10,000-40,000 years ago [1–5]. Some breed distinctions have been noted in archaeological and historical records, but most dog breeds have arisen within the past 150-200 years [6]. Increasingly intensive selective breeding over this comparatively short time has resulted in over 222 breeds [7] that vary in body shape, and in body size across two orders of magnitude [8,9]. This artificial selection pressure has been imposed to alter functional anatomy, behaviour, appearance, for specific working purposes or “desirable” companionable and aesthetic traits; some of which have more recently raised health and welfare concerns [10].

Within the musculoskeletal system of domestic dogs, variation in skull shape is well-studied (e.g. [11–13]), as are its genetic underpinnings [14], development [15], function [16,17], and implications for welfare [18,19]. Brachycephalic (short-faced) breeds, in particular, have received much attention, resulting in a wealth of research documenting shape, genetics, function, sociology and welfare, and leading to breed standard recommendations and a comprehensive understanding of the drivers and impacts of canine skull shape [20–22,18,23–26].

The postcranial musculoskeletal system, however, is less well-studied. Existing 3D geometric morphometric studies of limb elements across Carnivora are largely focused on locomotory adaptation or ecomorphology in an evolutionary context [27–30]. Consequently, we lack a comprehensive understanding of how intense artificial selection shapes postcranial form within a single species.

Domestic dogs present a unique opportunity to examine the extent, structure, and constraints of musculoskeletal diversification under artificial selection. Furthermore, certain breeds are known to develop specific disorders, but there is a relative paucity of the morphometric and functional data required to understand these associations. For example, which breeds are predisposed to hip dysplasia and patellar luxation are well-known from a clinical veterinary standpoint [31–33], but variation in the shape and functional anatomy of the appendicular skeleton is poorly understood. We do not currently possess sufficient data on breed-specific morphology and functional mechanics to determine the likely drivers of variation in postcranial musculoskeletal disease between breeds. In addition to aiding our understanding of welfare and disease risk, elucidating morphological and functional variation in domestic dogs presents the opportunity to investigate the evolution of canine postcranial morphology and biomechanics.

Existing 3D geometric morphometric studies of limb elements do not incorporate a large proportion of the shape of the bone; unambiguously homologous landmark positions are particularly limited along the shaft of long bones [34,29,30]. Some studies incorporate semilandmarks to increase the amount of shape captured [35], but the shape of the shaft and distal and proximal surfaces of the bone remain poorly sampled. Detecting subtle but functionally meaningful shape differences within a species requires both dense surface sampling and sufficiently large sample sizes. Landmark-based approaches restrict proportions of shape captured and may obscure fine-scale variation, thereby limiting the power to detect subtle but biologically meaningful variation within a species, where morphological differences may be smaller in magnitude than those observed in interspecific comparisons [36].

To overcome these limitations and enable large-scale, high-resolution quantification of appendicular morphology within domestic dogs, we used a landmark-free approach, Deterministic Atlas Analysis, to analyse scapula, humerus, radius, ulna, pelvic girdle, femur, tibia and fibula shape across 19 breeds (figure 1). This method is based on outputs from Deformetrica, software that produces a mean shape of an aligned and scaled set of meshes, applies a set of control points across the whole surface of the mean shape, and computes vector transformations required to transform that mean shape to each given mesh [37,38]. We ensured that these transformations are homologous by performing Procrustes alignment and scaling to a unit centroid size prior to analysis. Using this method, we found variation in the whole, holistic shape of each of the eight bones of interest. These findings constitute the first 3D analysis of limb bone shape between domestic dog breeds. Additionally, we investigated allometric effects, relative morphospace occupation, and modelled the relationships between shape, breed, breed category, age and body mass. We also investigated the degree and nature of integration between each of the elements, and explored the potential drivers of differences between breeds. By characterising whole-bone morphology and integration across breeds, this study establishes the extent and limits of appendicular shape diversity under artificial selection and provides a quantitative basis for linking postcranial morphology, function, and breed-specific disease risk.

**Figure 1.**
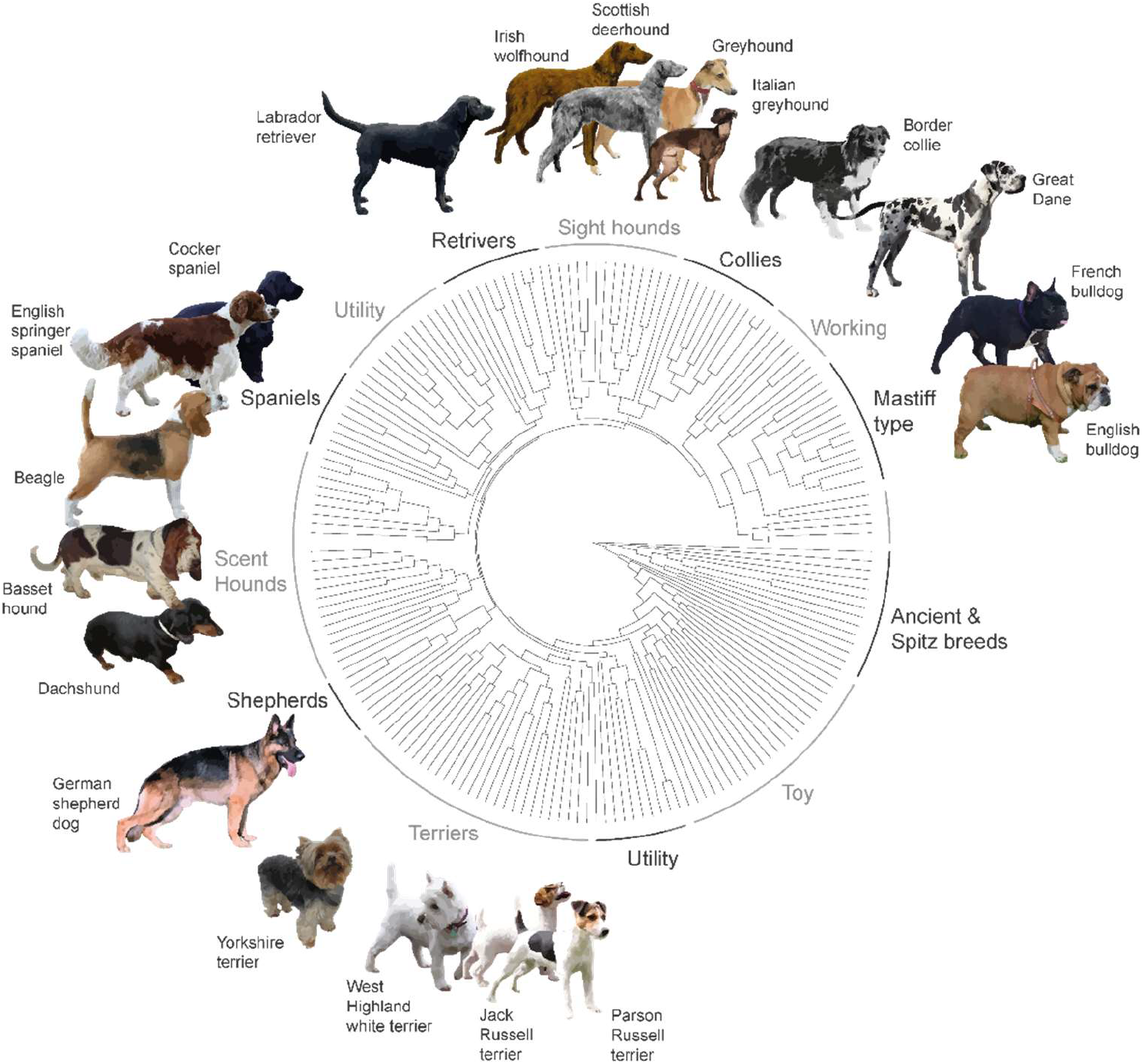
Dog breeds sampled in this study. Molecular phylogeny and breed groupings modified from Parker & Gilbert (2015). Images are modified from photographs accessed through Wikimedia Commons, all of which have been dedicated to the public domain under the Creative Commons CC0 1.0 Universal Public Domain Dedication.

## Methods

### Data Collection

Our dataset comprises 19 dog breeds (figure 1) and eight postcranial elements: scapula, humerus, radius, ulna, pelvic girdle, femur, tibia and fibula. Breeds were chosen to encompass a diverse range of body sizes, shapes and selective breeding factors i.e., different working types and companion animals. The dataset comprises 743 meshes of elements from 213 individuals, derived from computed tomography (CT) images of live dogs and surface scans of museum specimens. CT scans were produced by the Small Animal Teaching Hospital, University of Liverpool, accessed through the Liverpool Veterinary School Clinical Collaboration Platform (PACS system). We produced segmentations from CT data using SPROUT, a Python-based toolkit for semi-automated segmentation [39], and cleaned the resulting segmentations using Avizo [40], exporting them as .ply files. We produced surface scans in museum collections (table S1) using a Creaform Go!Scan 50, and processed raw scan data into .ply meshes using VXElements [41]. Museum specimens were included to increase temporal sampling.

Included CT specimens were scanned within the last 19 years (table S1), whereas museum specimens date between 2019 and 1891. Several museum specimens do not have exact dates attributed but are instead labelled, for example, “19th century” and are recorded as such (table S1). For each individual dog specimen, we segmented or scanned all elements that were available out of the eight postcranial elements that we aimed to sample. In order to analyse integration across limb elements for individual breeds, we aimed to produce at least five meshes for each breed. Museum specimens more commonly contained all eight elements of interest; however, in veterinary CT imaging, it is rare for whole dogs to be scanned at a sufficiently high resolution. More commonly, higher-resolution images appropriate for segmentation and 3D morphometrics contain only portions of the animal, usually individual whole limbs, or sections of the trunk or flanks that incidentally capture portions of the appendicular skeleton as part of a diagnostic investigation in veterinary clinical practice. Therefore, more than five individuals were typically required to fulfil our sampling goal for each bone.

For each scan from the Liverpool Veterinary School Clinical Collaboration Platform, we also recorded information associated with each individual at the time of scanning, including age, body mass, and incidence of relevant pathologies recorded in associated clinical notes. We recorded instances of the following pathologies: hip dysplasia, patellar luxation, carpal valgus, osteoarthritis, osteosarcoma, limb deformity, indeterminate lameness, and elbow incongruency/dysplasia. We coded individuals without any of these conditions as ‘none’ for modelling purposes. For each of the eight postcranial bone datasets, we re-assessed these categories, removing conditions irrelevant to that particular bone. For example, recorded osteosarcomas affecting the tibia were recorded as ‘none’ for the other elements e.g. femur.

### Shape analysis

We analysed shape using Deterministic Atlas Analysis, a landmark-free approach to shape analysis based on outputs of Deformetrica [37,38] analysed using established morphometry pipelines from [36,42]. Deformetrica takes a set of meshes and computes a mean shape or atlas, applies control points across the surface of the atlas and computes vector transformations from these control points to warp the mean shape to each mesh in the dataset. For analysis in Deformetrica, meshes must be watertight and aligned. We used Geomagic Wrap [43] (ver. 2017.0.2.18) to fill small holes using tangent fill, such that any filled space matches the curvature of the surrounding faces, and to decimate each mesh to 100,000 faces. To align the meshes, we applied a small number (5-7) of homologous landmarks in Stratovan Checkpoint [44] (figure S1, table S2). In the R statistical environment [45], we transformed these landmarks with Procrustes analysis and used *rotmesh*.*onto()* in *Morpho* [46] to align each mesh to its corresponding Procrustes coordinate position. To ensure that we are performing a true Procrustes transformation, we aligned the meshes first without scaling, and rescaled this set of meshes to a unit centroid size, as scaling using the internal function within *rotmesh*.*onto()* did not scale to unit centroid sizes across the set of meshes. We exported the meshes, converted them into .vtk format [47], and transferred them to a terminal using the Ubuntu operating system (ver. 22.04.4 LTS), as Deformetrica works only on Linux or Mac. Deformetrica requires users to input several model parameters; for each of the eight sets of meshes, we used a maximum of 150 iterations and a noise parameter of 1.0 and 20 timepoints. The number of timepoints represents how many iterations of warping are conducted between the atlas and each mesh [42]. The number of control points cannot be chosen directly and is instead determined by the input kernel width relative to the size of the atlas. Following Mulqueeney *et al*. [48] and Roberts *et al*. [36], we chose kernel widths such that each analysis used approximately 500 control points, requiring kernel widths between 0.426 and 1.75 mm for this dataset (table S3).

The outputs of Deformetrica include an atlas mesh (mean shape), the coordinates of the control points, 20 intermediate warps for each input mesh, and the momenta, which represent the vector transformations for each control point to each input mesh. Similar to a typical geometric morphometric analysis, the dimensionality of the raw outputs is reduced using principal components analysis (PCA) to aid interpretation, visualisation, and further analysis. However, the momenta are nonlinear data, requiring a slightly different approach. Following Toussaint *et al*. [42], we used kernel PCA (kPCA) [49] on the momenta to reduce the dimension to n, a chosen number of kernel principal components (kPCs). Here, we chose to calculate the same number of kPCs as a PCA would output in a traditional geometric morphometric analysis - the total number of specimens minus one.

### Allometry

To analyse and control for possible allometric shape variation, we extracted centroid sizes of the original meshes using the *cSize* function in Morpho, and logged them for further analysis. We ran multivariate regressions, regressing each kPC against log centroid size for each element. We extracted the residual kPC values for each kPC that was significantly correlated with log centroid size. These residual kPCs (kPC.resid) are the non-allometric portion of each kPC.

### Disparity

We quantified the volume of morphospace occupied by each breed for each element by taking the determinant of the covariance matrix for each group, which is proportional to the volume of the ellipsoid enclosing each group. We did this instead of calculating convex hull volumes to reduce the influence of outliers on volumes. To reduce the impact of differences in sampling between groups, we scaled ellipsoid volumes by taking the square root of the determinant, giving a linear scaling measure of relative morphospace volume.

### Integration analysis

We estimated integration between element pairs using two-block partial least squared analysis (PLS), using the *two*.*b*.*pls* function in geomorph. This function conducts PLS on two datasets. In PLS based on landmark data, the inputs would be arrays of Procrustes shape variables. Theoretically, the equivalent variables in Deterministic Atlas Analysis are the momenta, which represent the transformations from each control point on the mean shape required to warp to each mesh. However, the momenta are non-linear, meaning that these data are inappropriate for statistical analysis with linear methods, including *two*.*b*.*pls*. For this reason, following Roberts *et al*. [36], we ran integration analyses on the kernel principal components, which are a linearised representation of the shape data. Due to the nature of the datasets, we do not have all eight elements for the majority of specimens. Therefore, prior to each integration analysis, we created reduced kPC datasets containing only specimens for which we had both elements of interest, using the *intersect* function. Each dataset of paired elements in the integration analysis is therefore smaller than for any individual element. Several of these datasets, predominantly those including the tibia or fibula, contain fewer than 30 elements, necessitating cautious interpretation despite statistical significance.

### Modelling

To evaluate relationships between shape, breed, breed group, age, body mass and the presence of recorded appendicular pathologies, we ran a series of multivariate regressions of shape on these predictors. Age and body mass were logged and scaled prior to analysis. We used the *adonis2* function in the *vegan* package in R to perform non-parametric ANOVA (PERMANOVA) tests on the first six principal components for each element. We removed data for breeds with only one individual prior to analysis (e.g. English bulldogs and basset hounds). For each element, we ran three models: one assessing breed, one assessing breed group (e.g. working, hound), and one assessing breed given its inclusion within the relevant group (‘nested breed’). Breed groupings were assigned according to Parker and Gilbert [50]. We re-ran the three models using a “by = margin” term such that the function assesses the marginal effects of each term, so that all variables are considered simultaneously when assessing significance, and the potential significance of each predictor is tested individually. Following this, we investigated which pairs of breeds exhibit significant differences in bone shape, using the *pairwise*.*adonis2* function to assess shape differences between breeds, using (logged) age and (logged) body mass as covariates. All significance is evaluated at the p < 0.05 level.

## Results

### Shape

The first two kPCs cumulatively encompass 50-60% of total variation for each element (figure 2, figure 3). For most elements, there is substantial overlap in the morphospace of most breeds, with the exception of dachshunds and the single basset hound. These breeds instead occupy a distinct portion of the kPC1-2 morphospace for every element except for the pelvic girdle. Other than for the pelvic girdles, the maximum extent of kPC1 is occupied exclusively by dachshunds and the single basset hound in each element morphospace. Pelvic girdle shape overlaps for every breed in every dimension, except for French bulldogs which show a degree of separation across kPCs 2, 3 and 4 (figure 3A, figure S3A). For the scapula, dachshunds and the basset hound separate exclusively across kPC2. Dachshunds and the basset hound are most distinctly separated from every other analysed breed in the tibia and fibula morphospaces. In addition to the distinct separation of dachshunds from every other breed, the proportion of the morphospace occupied by dachshunds is much larger than that of every other breed for every element except for the pelvic girdle and fibula.

**Figure 2.**
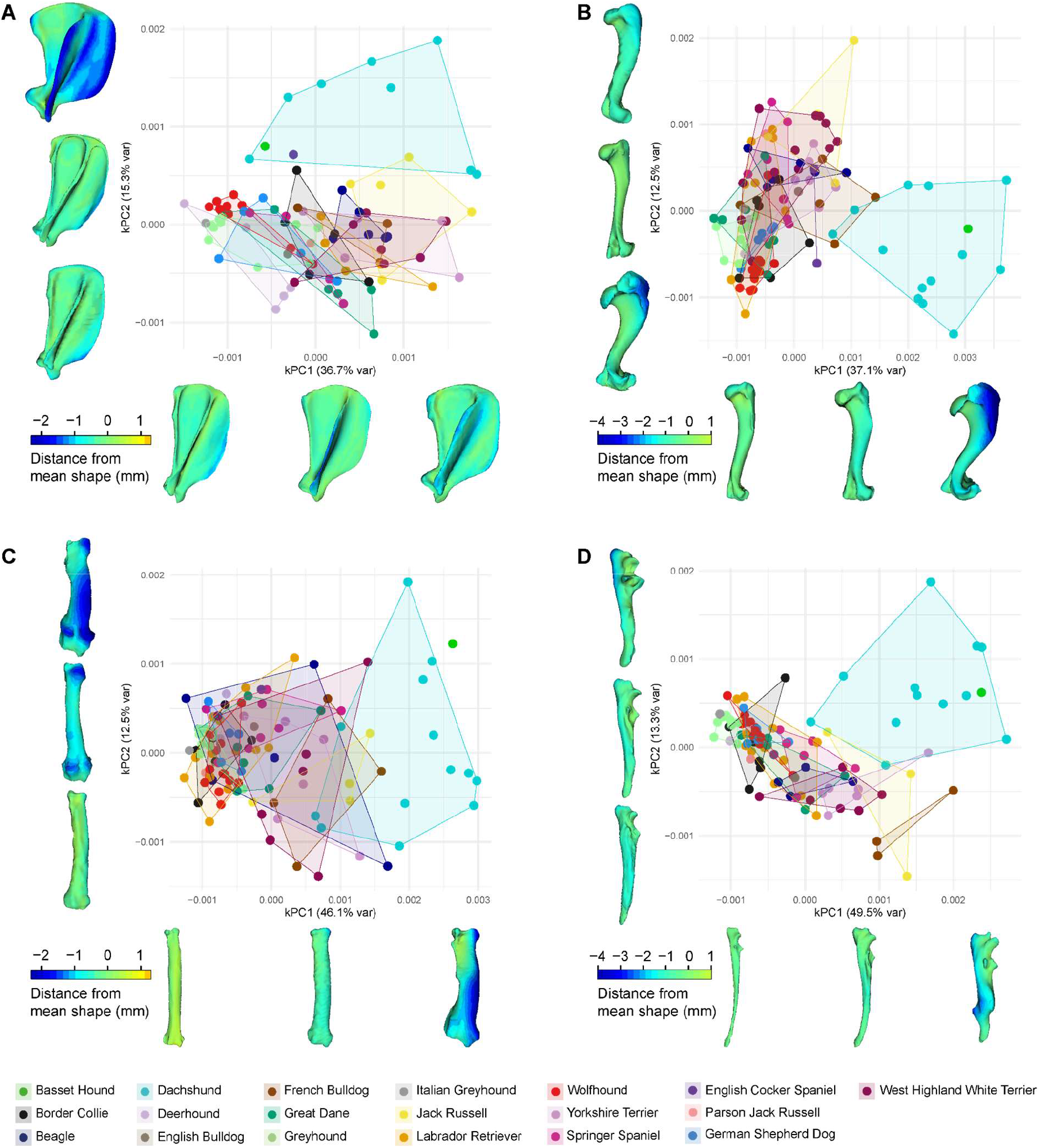
Shape variation in forelimb elements represented by the first two kernel principal components (kPCs) for the scapula (**A**), humerus (**B**), radius (**C**) and ulna (**D**) alongside meshes of elements at minimum, median and maximum values of kPC1 and 2 coloured by distance from the mean shape of that element.

**Figure 3.**
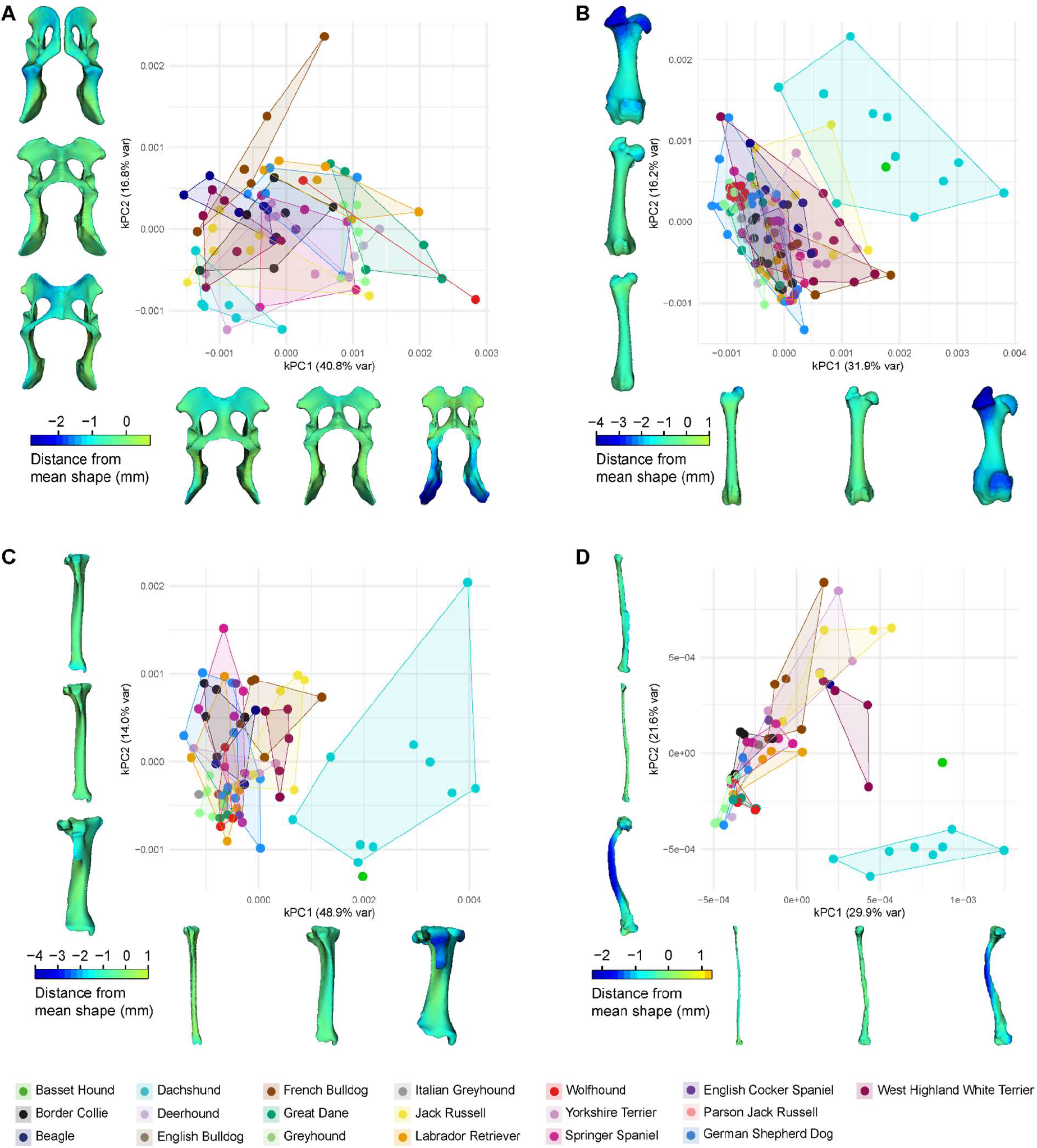
Shape variation in forelimb elements represented by the first two kernel principal components (kPCs) for the pelvic girdle (**A**), femur (**B**), tibia (**C**) and fibula (**D**) alongside meshes of elements at minimum, median and maximum values of kPC1 and 2 coloured by distance from the mean shape of that element.

Ellipsoid volume, a measure of the amount of morphospace occupied by each breed, is highest for most elements in dachshunds. For the humerus, radius, ulna, femur and tibia, dachshunds occupy 1-2 orders of magnitude more of the morphospace than the other breeds (table S4), whereas the scapulae, pelvic girdles and fibulae occupy moderate to low volumes. The largest portions of the morphospaces for the pelvic girdles and fibulae are occupied by Yorkshire terriers, and Great Danes occupy the largest area of scapular morphospace. Greyhounds, wolfhounds and Border collies consistently occupy small areas of each morphospace.

For each element, kPC1 encompasses variation in overall aspect ratio, except for the scapulae and pelvic girdles, for which aspect ratio varies across kPC2 (figure 2, figure 3). For all elements except for the girdles, low values of kPC1 represent proximodistally elongate, mediolaterally and/or craniocaudally restricted (i.e., elongate and relatively narrow) morphologies, common to Labrador retrievers, wolfhounds, greyhounds, German shepherd dogs and Great Danes. High values of kPC1 represent relatively thickset morphologies that are proximodistally compact and mediolaterally and/or craniocaudally expanded, common to dachshunds, West Highland white terriers and Jack Russell terriers. The shaft of each element represents a greater proportion of the total length at maximum values of kPC1. kPC2 represents more subtle shape changes. For some elements, kPC2 represents the presence or absence of curvature. For example, fibulae at high kPC2 are straight, aligned with the proximodistal axis, whereas fibulae at minimal values of kPC2 deviate from alignment with this axis due to strong lateral curvature (figure 3D).

Dachshunds represent the extreme expression of this morphology. At high values of kPC2, the caudal margin of the scapula is strongly curved, compared to a relatively straight profile at low values (figure 2A). For the humerus, radius, ulna, femur and tibia, kPC2 encompasses differences in the shape of proximal and distal processes. For example, humeri at the minimum extent of kPC2 have more prominent greater and lesser tubercles, and the medial margin of the caudal aspect of the trochlea is more angular. For femora at the positive end of kPC2, the processes of the proximal end, including the greater tubercle, are more pronounced, and the trochlear grooves on the distal end are deeper (figure 3B).

### Allometry

Accounting for allometry by plotting residuals against log centroid size for kPCs with a significant allometric relationship, has little effect on the structure of kPCs other than kPC1 (figure S4,S5). Despite significant (p < 0.05) relationships between several kPCs and log centroid size for all individual elements, the fit and degree of correlation is low (table S5). The residuals of kPC1 (kPC1.resid), however, do differ from the original values of kPC1. In all further analyses herein, we use the residual values of each kPC for which the regression with centroid size is significant. Constructing morphospaces with kPC1.resid results in almost complete overlap of every breed except for dachshunds and basset hounds. There is also increased overlap of dachshund convex hulls with other breeds in each residual morphospace except for the pelvic girdle (figure S4,S5), and the size of the convex hulls representing dachshunds is either the same or larger when allometry is accounted for within this analysis.

### Correlates of shape variation: breed, age, body mass and pathology

#### Scapula

Age is not a significant predictor of scapula shape across dog breeds (table S7). Body mass is a significant, albeit very marginal, predictor of scapula shape; the analyses that assess shape across breed and breed category independently indicate that (logged) body mass accounts for ∼3-5% of shape variation. Breed, breed group, and nested breed are all significant predictors. In the breed-only analysis, breed explains the largest proportion of shape (R^2^ = 0.40). Breed group differences are significant but small; breed group alone explains ∼18% of variance, and nested breed explains 21%. Scapula shape is significantly and strongly distinct (R^2^ ≥ 0.45) between sixteen breed pairings once the small effect of body mass is accounted for (table S8). Beagles and Labrador retrievers appear most frequently in these pairs (six and five pairs respectively), but the largest proportion of shape variation attributable to breed differences is modelled between Great Danes and deerhounds (74%), greyhounds (68%) and Yorkshire terriers (67%).

#### Humerus

Breed, breed group, and nested breed are significant predictors of humerus shape (table S7). Breed alone explains the largest proportion of shape (R^2^ = 0.34). Breed group and nested breed are significant but explain a lower proportion of shape variation (R^2^ = 0.17). Neither age nor body mass are significantly associated with humerus shape when breed is considered. However, in the breed-group-only model, body mass is recovered as a significant but marginal predictor, accounting for just 4% of total variation in shape. Dachshunds show significant and strong differences in humerus shape (R^2^ = 0.4-0.5) in pairwise comparisons with eight other breeds (table S8). English springer spaniel humeral shape is significantly different to Border collies and Labrador retrievers (R^2^ = 0.54).

#### Radius

We recover significant differences in radius shape globally between breeds, breed groups and nested breed (table S7). Radius shape is significantly but marginally (R^2^ = 0.06-0.07) associated with age, and is not significantly associated with body mass. In the pairwise comparisons, we recover strong significant differences in radius shape between Great Danes and French bulldogs, between Labrador retrievers and four other breeds, and between dachshunds and eight other breeds (table S8).

#### Ulna

Ulna shape is significantly different between breeds, breed groups, nested breeds, and is significantly associated with both age and body mass (table S7). In the breed-only model, breed accounts for 32% of the observed variation in ulna shape. In the breed-group model, group accounts for 24%, but in the nested breed model, group accounts for only 8%. Age and body mass account for 4-8% of the total variation in ulnar shape (figure S8,S9). Pairwise comparisons demonstrate significant and strong differentiation between 20 breed pairs (table S8). Dachshunds separate significantly and clearly from every other breed in this multivariate Euclidean space (R^2^ = 0.5-0.7). French bulldogs also separate from every other breed except for Yorkshire terriers and Jack Russell terriers.

#### Pelvic girdle

Pelvic girdle shape varies significantly between breeds, breed groups and nested breed (table S7). The proportion of variance attributed to breed/group is lower than that of other examined elements; in the breed-only model, breed accounts for 29% shape variation, in the group-only model, breed group accounts for 16%, and in the nested model, breed accounts for just 13%. Age has a small but significant association with pelvic girdle shape, accounting for 3-7% of the total variation. When the breed category is taken into account, body mass is insignificant, but in the breed group only model, body mass accounts for 3% of the total variation (figure S8). Pelvic girdle shape is significantly different between 36 breed pairs (table S8), more than any other examined element. Pelvic girdle shape is significantly distinct between Jack Russell terriers and every other breed except for English springer spaniels, beagles and Irish wolfhounds; dachshunds are similarly separated from every other breed except for these three breeds and Border collies.

#### Femur

Femur shape is significantly different between breeds, breed groups, and nested breeds (table S7). Similar to other examined limb elements, when breed is examined in isolation from breed group it accounts for a higher proportion of variation (31%), compared to 16% for nested breed. When modelled without the effect of breed, breed group accounts for 15% total variation. In the breed group only and nested breed models, body mass has a statistically significant but marginal impact on femur shape, accounting for just 1-3% of the total variation. In the pairwise PERMANOVA, we find that Jack Russell terrier femur shape is significantly and strongly (R^2^ ≥ 0.45) different compared with nine of the other breeds, i.e., all other breeds in the model except for dachshunds, West Highland white terriers and German shepherd dogs (table S8).

#### Tibia

Breed is the dominant factor affecting tibia shape (table S7). In their respective models, breed and breed group are significant predictors of tibia shape; breed alone explains the largest proportion of shape (R^2^ = 0.45), whereas breed group alone explains ∼39% of variance. In the breed group model, body mass is significantly but marginally associated with shape, explaining just 5% of variation. Significant and strong differentiation is modelled between 18 breed pairs (R^2^ = 0.5-0.7). Eight of these pairs include dachshunds (table S8).

#### Fibula

Fibula shape varies significantly between breeds, breed groups and nested breeds (table S7). In the breed-only model, breed accounts for 47% variation, and in the breed group model, group accounts for 39%. Therefore, breed in the nested model accounts for only ∼8% total variation. In all three models, body mass is not recovered as a significant factor, but age is a significant, marginal factor, accounting for 4-5% total variation. In the pairwise PERMANOVA, the only significant pairwise differences in shape are recovered between dachshunds and seven other breeds (table S8).

### Pathology

Across the whole dataset, we recover weak relationships between bone shape and limb pathologies as recorded by veterinarians (figure S7). We only recover significant relationships between shape and presence of pathology for the radius (p = 0.04, R^2^ = 0.1) and humerus (p = 0.03, R^2^ = 0.1). Subsequent pairwise analysis demonstrates that this relationship recovered for radii is driven by a significant difference in radius shape between dogs with identified elbow incongruency and those designated as ‘normal’ (i.e., with no recorded appendicular pathologies) (table S9). For the humerus, we also recover significant differences between ‘normal’ individuals and those with recorded elbow incongruency, and between ‘normal’ dogs and those with carpal valgus. The statistical relationship between shape and pathology is maintained when breed is taken into account for the radius (p = 0.04, R^2^ = 0.1), but not the humerus (p > 0.05) (Table S7)

Variation in the shape of each bone across the whole dataset is dominated by breed differences. Thus, we ran additional models within each breed where possible (n ≥ 5). For most elements, we do not recover any significant relationships (table S10). We do, however, recover significant relationships between shape and the presence of appendicular pathologies for the pelvic girdle in Yorkshire terriers (p = 0.045, R^2^ = 0.35), and Jack Russell terriers (p = 0.046, R^2^ = 0.51), and the radius in West Highland white terriers (p = 0.047, R^2^ = 0.50) and Labrador retrievers (p = 0.046, R^2^ = 0.55).

### Integration

Two-block partial least squares analyses demonstrate pervasive and predominantly positive covariation among limb elements across breeds, indicating strong morphological integration within both forelimb and hindlimb. The magnitude of integration is consistently high across most element combinations (figure 4, figure S6) and statistically significant for all pairings except pelvic girdle-fibula (table S6). In the forelimb, integration is uniformly strong across all element pairings, with particularly tight associations between adjacent elements (humerus-radius, humerus-ulna, and radius-ulna). Scapular shape also covaries strongly with more distal elements, indicating coordinated variation across the entire forelimb module. In the hindlimb, integration is strongest proximally (pelvic girdle-femur; femur-tibia), whereas pairings involving the fibula show greater dispersion and increased breed-specific variability. Although distal hindlimb elements remain positively integrated, their covariation is less consistent than that observed in the forelimb.

**Figure 4.**
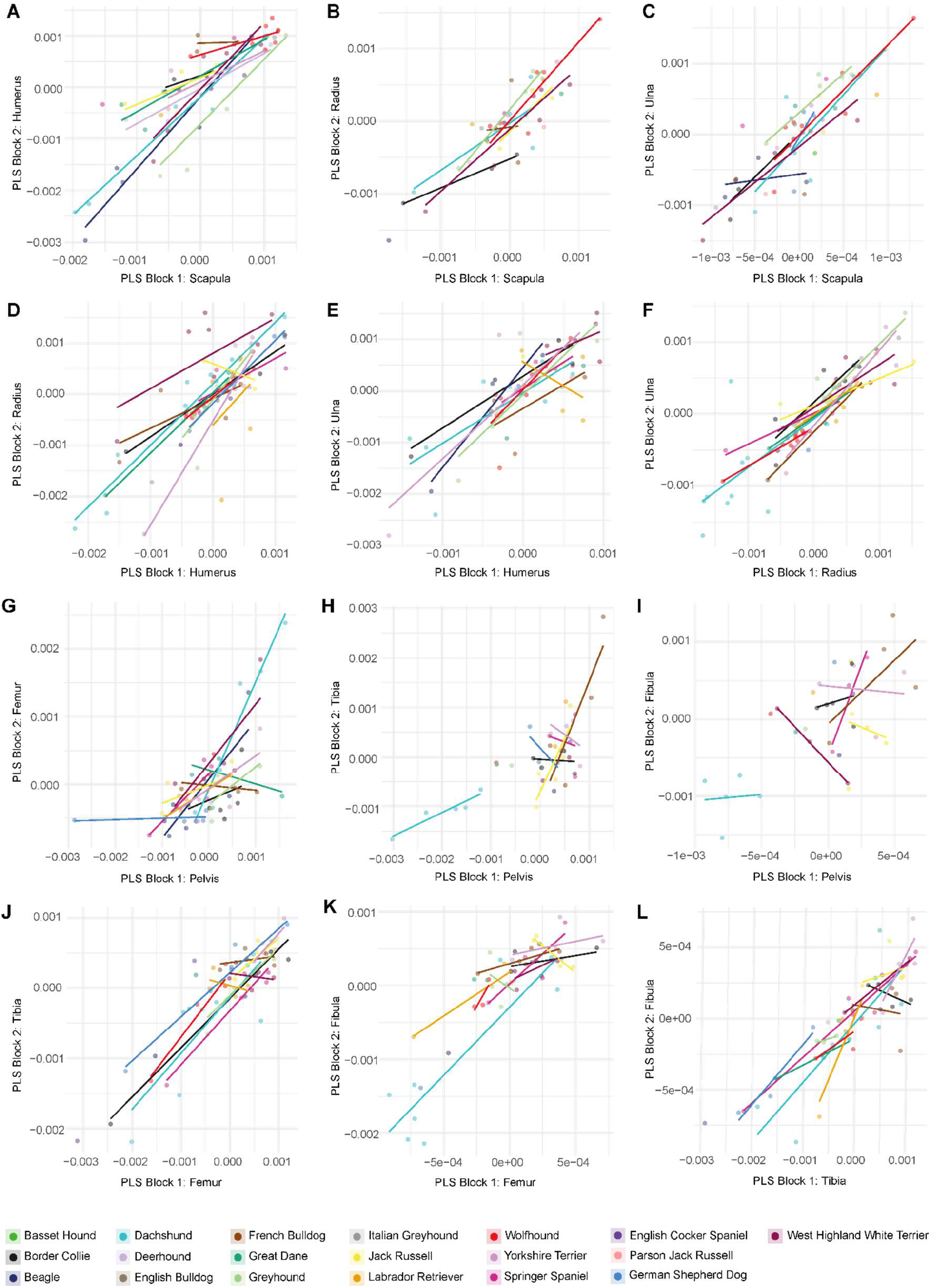
Integration of limb elements: Partial least squares (PLS) analysis separated by breed **A** Scapula vs humerus **B** Scapula vs radius **C** Scapula vs ulna **D** Humerus vs radius **E** Humerus vs ulna **F** Radius vs ulna **G** Pelvic girdle vs femur **H** Pelvic girdle vs tibia **I** Pelvic girdle vs Fibula **J** Femur vs tibia **K** Femur vs fibula **L** Tibia vs fibula.

Partitioning PLS analyses by breed reveals that forelimb elements are strongly integrated in all sampled breeds, with largely parallel breed-specific slopes indicating conserved directions of covariation despite differences in magnitude (figure 4). A notable exception is the humerus-ulna relationship in Labrador retrievers, where the direction of covariation deviates from that observed in other breeds. In contrast, hindlimb integration exhibits greater inter-breed heterogeneity. The tibia and fibula are strongly integrated in most breeds but show reduced or inconsistent covariation in Border collies and French bulldogs.

Similarly, femur-tibia covariation appears more variable in French bulldogs, Labrador retrievers, and West Highland white terriers relative to other breeds, although overall integration remains positive. Pelvic girdle-femur covariation follows a similar trajectory in most breeds; however, in German shepherd dogs and French bulldogs, pelvic girdle-femur covariation exhibits a lower apparent magnitude compared to other breeds.

Several breeds occupy distinct regions of hindlimb PLS morphospace, most notably dachshunds in pelvic girdle-tibia, pelvic girdle-fibula, and femur-fibula comparisons, and German shepherd dogs in pelvic girdle-femur. This indicates displacement along the primary axis of covariation rather than alteration of the covariance structure. These patterns suggest that breed-specific hindlimb morphologies largely reflect divergence along conserved covariance trajectories, with localised reductions in integration in certain comparisons.

## Discussion

Despite the diversity of body size and overall body shape across dog breeds, the limb and girdle bones are remarkably similar in shape in most breeds studied here. Extreme variation in the shape of the canine skull is well-documented [11,15,12,51,13,16], but our results suggest that, amongst most breeds, there is much less variation in shape within the postcranial skeleton. Accounting for allometry further reduces the observed variation in each elemental morphospace. For example, in the non-allometry-corrected kPC1-2 morphospace for the femur, the convex hulls for most breeds overlap markedly, although the convex hulls representing Jack Russell terriers, Yorkshire terriers and French bulldogs separate slightly, and dachshunds separate further (figure 3B). Once allometry is accounted for, dachshunds remain separated, but these other small breeds overlap almost entirely with every other breed (figure S5B). Thus, the primary axis of variation across elements is largely allometric. Dachshund limb bone and scapula shape deviate prominently from all other breeds, even when the allometric component of shape is removed, characterised by an overall relative robustness, increased curvature, and exaggerated processes on the joint surfaces (figure 2, figure 3, figure S4, figure S5).

Further to strong separation from other breeds within the morphospace, the amount of morphospace occupied by dachshunds is one to two orders of magnitude larger than other breeds for most elements (table S4). This large shape disparity cannot be explained by age or body mass (figure S8, figure S9, table S7). A potential source of variation within this dataset is the inclusion of both recent and historical data. Most sampled dachshunds are veterinary specimens CT scanned within the past 20 years as part of routine veterinary practice for the purposes of diagnostic investigation for clinical conditions with full owner consent and ethical permission. Three sampled dachshunds are from museum collections. Two collected in 1950, one of which is in all of the bone samples except for the fibula, and the other of which is in all samples except for the fibula and pelvic girdle, and an additional specimen collected in the late 1800s (exact date unknown), which is included in all samples except for the fibula and pelvic girdle. The oldest specimen sits within the portion of the morphospace occupied by many other breeds, and each individual element is visibly more similar to modern dogs of other breeds than to recent dachshund specimens (figure S10). The specimens from 1950 sit closer towards modern dachshunds. Despite this temporal variation in the dataset, this also does not explain the volume of morphospace occupation; the largest spread in each morphospace is made up of modern specimens.

Another possible explanation for the large volume of morphospace occupied by dachshunds is pathology. Certain clinical conditions may impact bone shape; for example, dogs with hip dysplasia may have consistently different femoral or pelvic girdle shapes compared with individuals that do not have the condition. However, any relationships we recover between shape and pathology explain only a marginal proportion of variation across the global dataset, and we do not recover significant differences within the dachshund-only subsets (figure S7, table S7, table S10). It is possible that larger datasets focused principally on dachshunds with different appendicular pathologies may recover some relationship, but within the scope of this study, we do not recover any such association. Another potential source of extreme shape variation within dachshunds is the biomechanical impact on bone shape during ontogeny, which potentially differs in dachshunds due to their unique body shape. Type and amount of exercise have been linked to the occurrence of intervertebral disc calcification and intervertebral disc disease in dachshunds [52,53]. Differences in the biomechanical impact of exercise and ontogeny in dachshunds may also impact the appendicular skeleton to a greater degree than breeds that have less extreme limb morphologies. Testing this is beyond the scope of the data collected for this study, but presents a viable future direction for research on dachshund health and welfare.

Dachshunds are a chondrodysplastic breed; they have a form of canine dwarfism. Genetic research has discovered a retrogene associated with a short-legged phenotype in dogs [54], therefore we are potentially recovering a difference in limb bone shape between breeds with chondrodysplasia and those without these genetic markers. However, studies have found the FGF4 retrogene responsible for this condition in multiple breeds included in this study, such as: dachshunds, basset hounds, West Highland white terriers, beagles, and Yorkshire terriers (at a lower frequency) [54–56]. We do not find the same limb bone shape in all of these breeds; once allometry is corrected for, limb bone shape in West Highland white terriers, beagles and Yorkshire terriers differs little from other breeds. This genetic variation likely contributes to some shape variation but cannot alone explain the extreme morphology observed in dachshunds.

Genetic variation within dachshunds may contribute to the relatively high degree of shape variation within the breed. Standard dachshunds include smooth-haired, long-haired and wire-haired varieties. Smooth-haired standards are the oldest variety, then wire and long-haired varieties were developed by cross-breeding with various other breeds, resulting in populations with genetic distinctions [57–59]. Coat varieties have been linked to genetic variation that can result in musculoskeletal disease such as osteogenesis imperfecta, which is a congenital disease that affects the development of connective tissue, and has been found almost exclusively in wire-haired dachshunds [60]. We do not have data on the coat type of dachshunds included in this study, but the inclusion of different coat varieties within the dachshund dataset may be a contributory factor to the large morphological spread of limb bone shape.

Since 1977, under the UK Kennel Club system, cross-coat matings are not permitted; however, the registered variety follows “the breed to which its coat most closely conforms” [61] regardless of the coat type of the parents. The longhair gene is recessive to both wire and smooth hair genes, and the wire hair gene is dominant to smooth. The breeding history of dachshunds has created incompletely separate genetic groups that may contribute to variation in bone shape across the breed recovered herein.

Furthermore, dogs bred under different breed standards may develop different bone shapes, the majority of this dataset samples dogs in the United Kingdom, but also includes individuals from the US. The provenance of each individual beyond the country they were scanned or accessioned in is not known, however variation found herein across dachshunds highlights the potential for future studies investigating variation within the breed, across coat types, size categories, countries of origin and life histories including exercise levels.

Despite pronounced deviations in limb proportions, the magnitude of integration within the appendicular skeleton of dachshunds remains consistently high. In the forelimb, covariation is broadly similar across all element combinations and across breeds, with largely conserved directions of integration. Some breed-specific differences are evident. For example, although the humerus and ulna in Labrador retrievers are as strongly integrated as in other breeds, the direction of covariation differs, suggesting a modification in the scaling relationship rather than a reduction in integration (figure 4E). Variation among breeds is more apparent in the hindlimb. Dachshunds differ markedly from other breeds in the shapes of the femur, tibia, and fibula, whereas pelvic girdle morphology is comparatively conserved. However, rather than indicating a change in covariance, integration analysis shows that dachshunds are displaced along the primary axis of covariation shared with other breeds. Thus, integration between the pelvic girdle and distal hindlimb elements appears to be maintained, with selective modification occurring along established trajectories of morphological integration. Similar displacement patterns are observed in other hindlimb comparisons, suggesting that extreme breed morphologies reflect divergence along conserved covariance axes rather than disruption of underlying modular relationships. Consistently high degrees of integration within the appendicular skeleton are unsurprising, as a loss of coordinated covariation within or between limb segments would likely compromise locomotor function. Instead, the results indicate that artificial selection on limb proportions operates within a conserved framework of developmental and functional integration.

Herein, we find significant differences in the shape of limb elements between domestic dog breeds, but also that the majority of this shape variation is allometric. The first kernel principal component represents an allometric gradient in aspect ratio. In larger breeds, the limb elements are elongated; in smaller breeds, the shafts of limb elements are proportionally shorter, and the proximal and distal portions are relatively robust. We further find that, beyond allometry, the dominant explanatory variable of shape across kPC1-6 is breed. Body mass and age, when significant, are only marginally associated with bone shape when breed is accounted for, and the presence of limb-related pathologies only has moderate relationships to pelvic girdle shape in smaller breeds. However, we did not sample specifically for pathologies, so further work is needed to clarify this relationship. Dachshunds in particular dominate each morphospace, separating distinctly from other breeds even when allometric effects are removed. The sole outlier to dachshund exceptionalism is the pelvic girdle, which exhibits comparatively limited divergence across breeds. This relative conservatism may reflect functional constraints, potentially including reproductive demands.

Consistent with functional constraints on the appendicular skeleton, integration between elements is high despite breed variation. Within-breed shape variation is relatively large in dachshunds, again except for the pelvic girdle. We find no statistical link to body mass variation, age, or pathology within dachshunds. Some of the variation is explained by the distribution in specimen age - the elements of individuals from the 19th and mid 20th century are more similar in shape to non-dachshund breeds. However, most variation is within modern dachshunds, potentially related to the inclusion of multiple genetic lineages within standard dachshunds. Here we demonstrate the importance of quantifying phenotype for mapping of potential genotypes; several of the breeds included herein, including beagles, Yorkshire terriers, Jack Russell terriers and French bulldogs, have the genetic markers of chondrodystrophy, but only dachshunds and basset hounds display extreme morphologies of the appendicular skeleton. This underscores the importance of phenotypic quantification for interpreting underlying genetic architecture.

Overall, our findings show that even within a highly integrated skeletal system, selective pressure and genetic diversity can drive extreme, breed-specific morphologies. Quantitative analyses of postcranial variation provide a framework for linking phenotype to genotype, assessing breed health, and informing responsible breeding practices. Extending this framework to incorporate muscle architecture and joint mechanics will be essential for determining how observed shape differences translate into functional performance and disease susceptibility. The variation in bone morphology quantified here may alter the geometry of muscle attachment areas and thereby influence musculoskeletal kinetics. Given the strong integration observed across the appendicular skeleton, such effects are likely to extend beyond individual elements to the wider postcranial skeleton, including the vertebral column. In breeds exhibiting extreme morphologies, alterations to the vertebral column may be similarly pronounced. Dachshunds, for example, are predisposed to intervertebral disc disease [62], but the causes of this breed-specific susceptibility remain poorly understood. This study underscores the necessity of adopting a whole-organism perspective when assessing breed standards and their biological impacts, and establishes a quantitative foundation for linking skeletal variation to functional and clinical outcomes that support evidence-based improvements in canine health and welfare.

## Acknowledgements

This work was supported by Biotechnology and Biological Sciences Research Council grant BB/X014819/1

We would like to thank T Maddox for CT scan access and R Portela Miguez, A Farrel, K Zyskowski and T Hsu for specimen access. We would like to thank M Camaiti for the useful discussion.

## Ethics

Ethics approval for the use of CT scans was granted by the University of Liverpool Veterinary Research Ethics Committee (VREC), code VREC1311 for all non-greyhound breeds and VREC1339 for greyhounds.

## Data accessibility

Code used to generate these results is available through github https://github.com/LucyEmmaRoberts/landmark-free

Meshes, landmarks: Dryad repository https://doi.org/10.5061/dryad.fttdz0978 (repository will be publicly available on publication)

All other data supporting this article is available as part of the supplementary material.

