## Supplementary figures and tables for "Extreme disparity in the appendicular skeleton of domestic dogs (*Canis familiaris*)"

### Supplementary Information for: Extreme disparity in the appendicular skeleton of domestic dogs (*Canis familiaris*)

This supplementary information file contains:

Supplementary figures 1 to 10

Supplementary tables (or table captions) 1 to 10

Supplementary Figure 1: Landmarks used to align meshes alongside principal component 1 and 2 plots generated from geometric morphometric analysis of the corresponding landmarks. **A** Scapula **B** Humerus **C** Radius **D** Ulna **E** Pelvic girdle **F** Femur **G** Tibia **H** Fibula

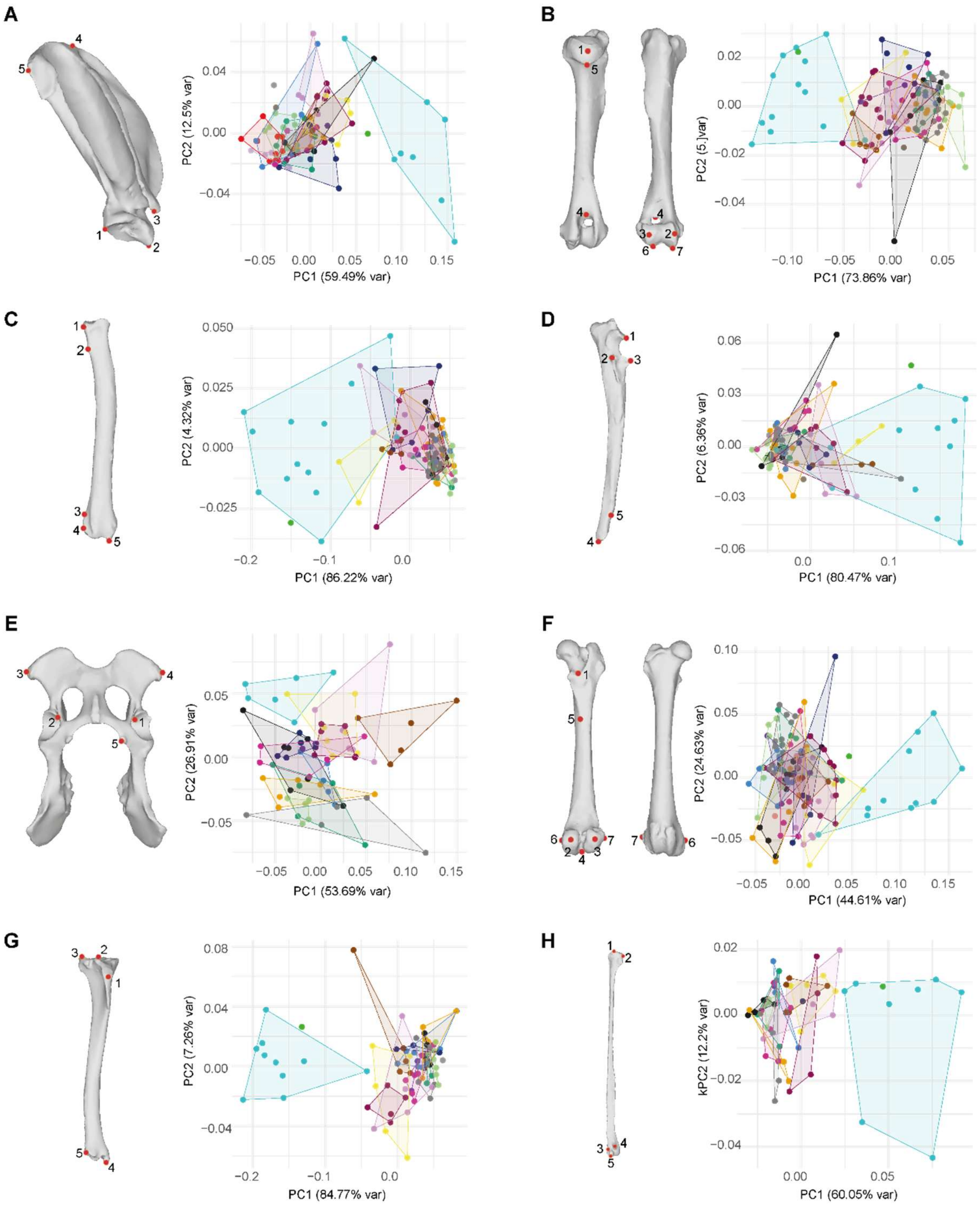

Supplementary figure 2: Kernel Principal Components (kPC) 1-6 Forelimb elements **A** Scapula **B** Humerus **C** Radius **D** Ulna

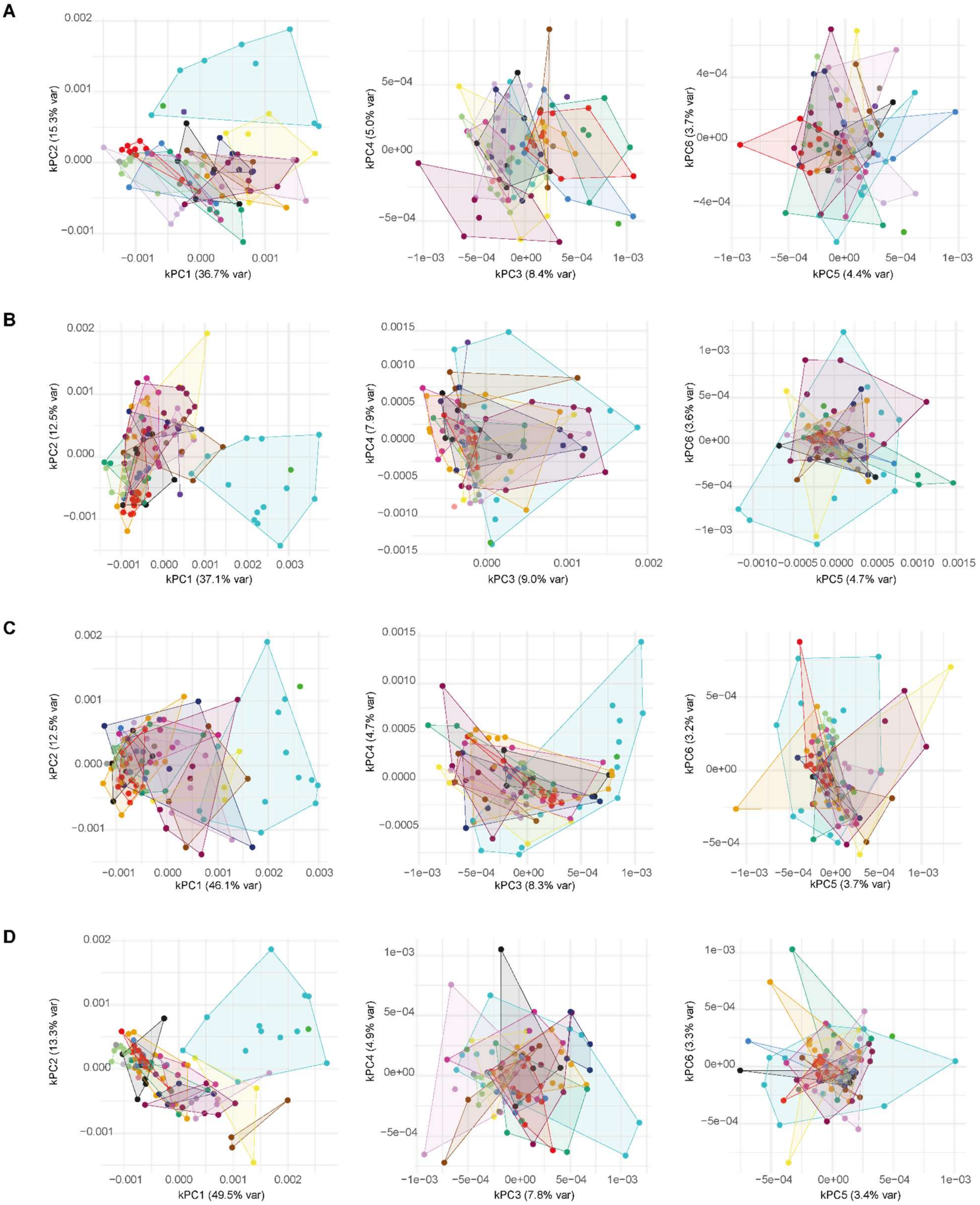

Supplementary figure 3: Kernel Principal Components (kPC) 1-6: Hindlimb elements **A** Pelvic girdle **B** Femur **C** Tibia **D** Fibula

**A**

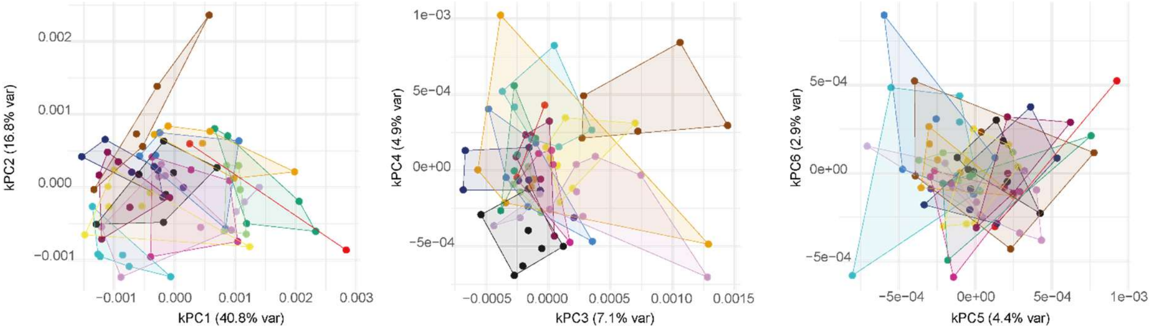


**B**

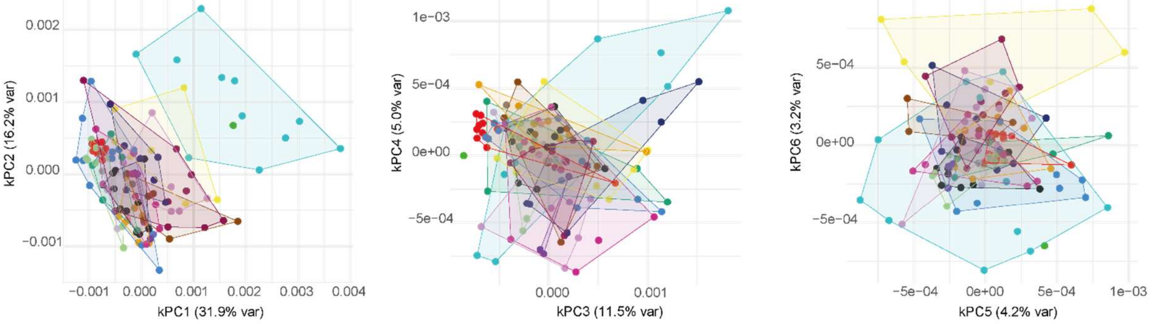


**C**

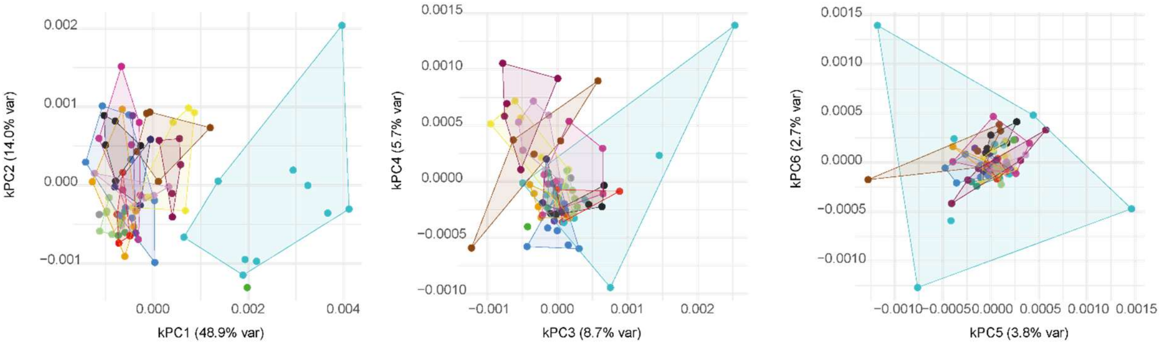


**D**

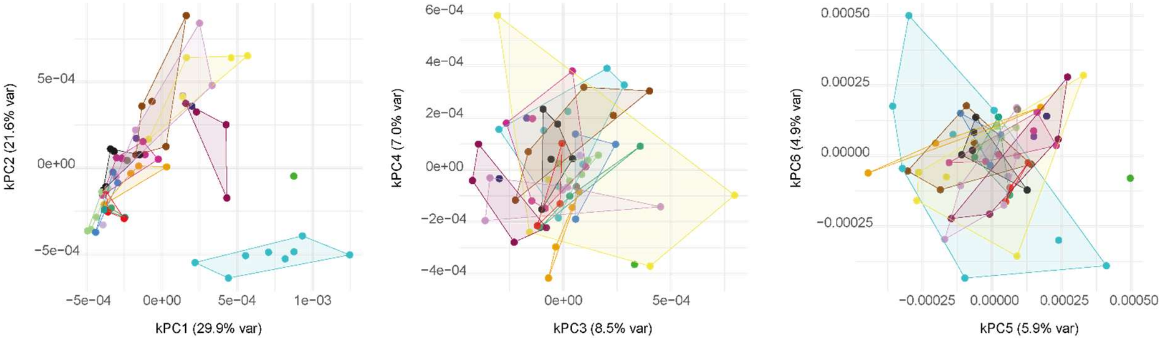

Supplementary figure 4: Kernel Principal Components (kPC) 1-6 corrected for allometry where necessary: Forelimb elements **A** Scapula **B** Humerus **C** Radius **D** Ulna

**A**

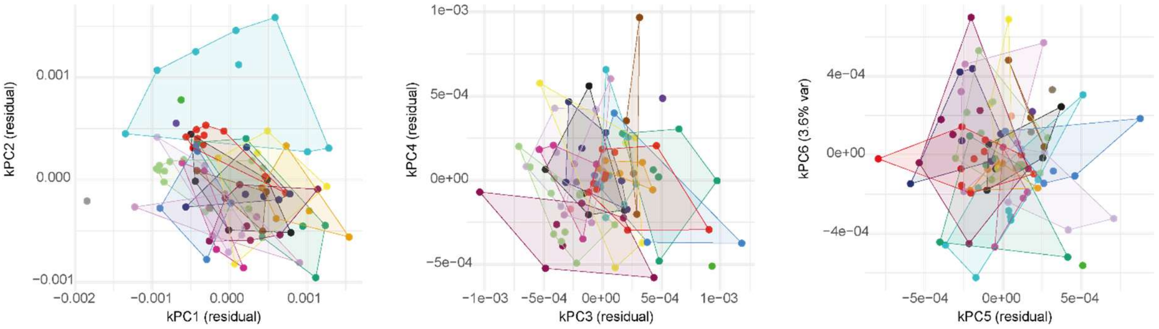


**B**

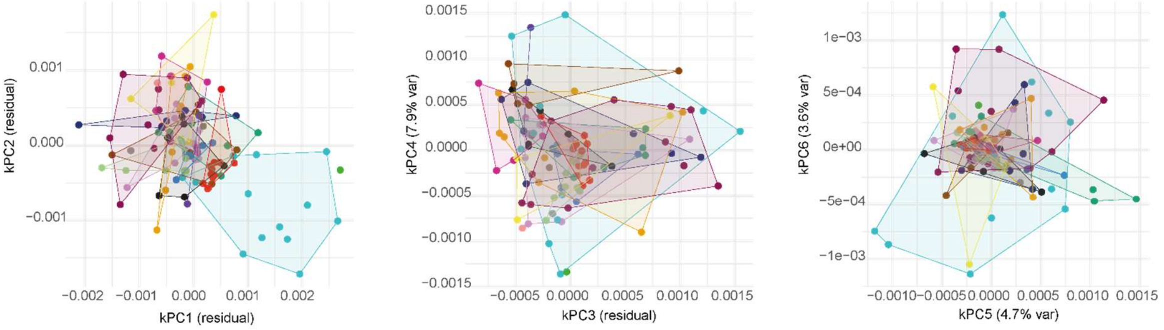


**C**

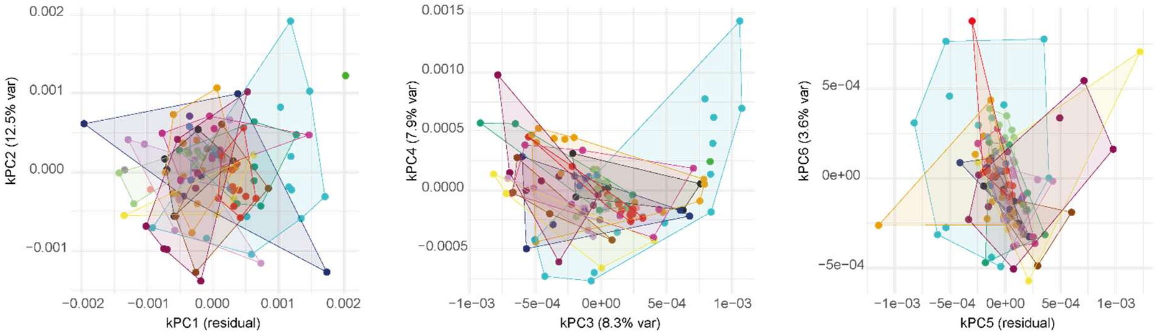


**D**

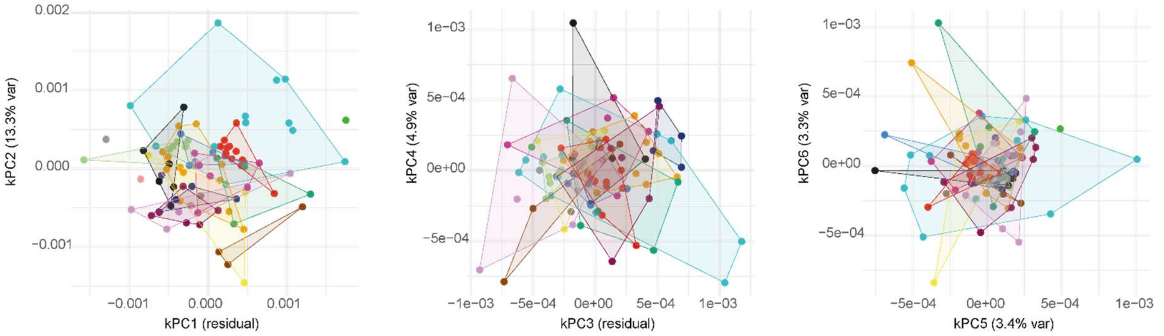

Supplementary figure 5: Kernel Principal Components (kPC) 1-6 corrected for allometry where necessary: Hindlimb elements **A** Pelvic girdle **B** Femur **C** Tibia **D** Fibula

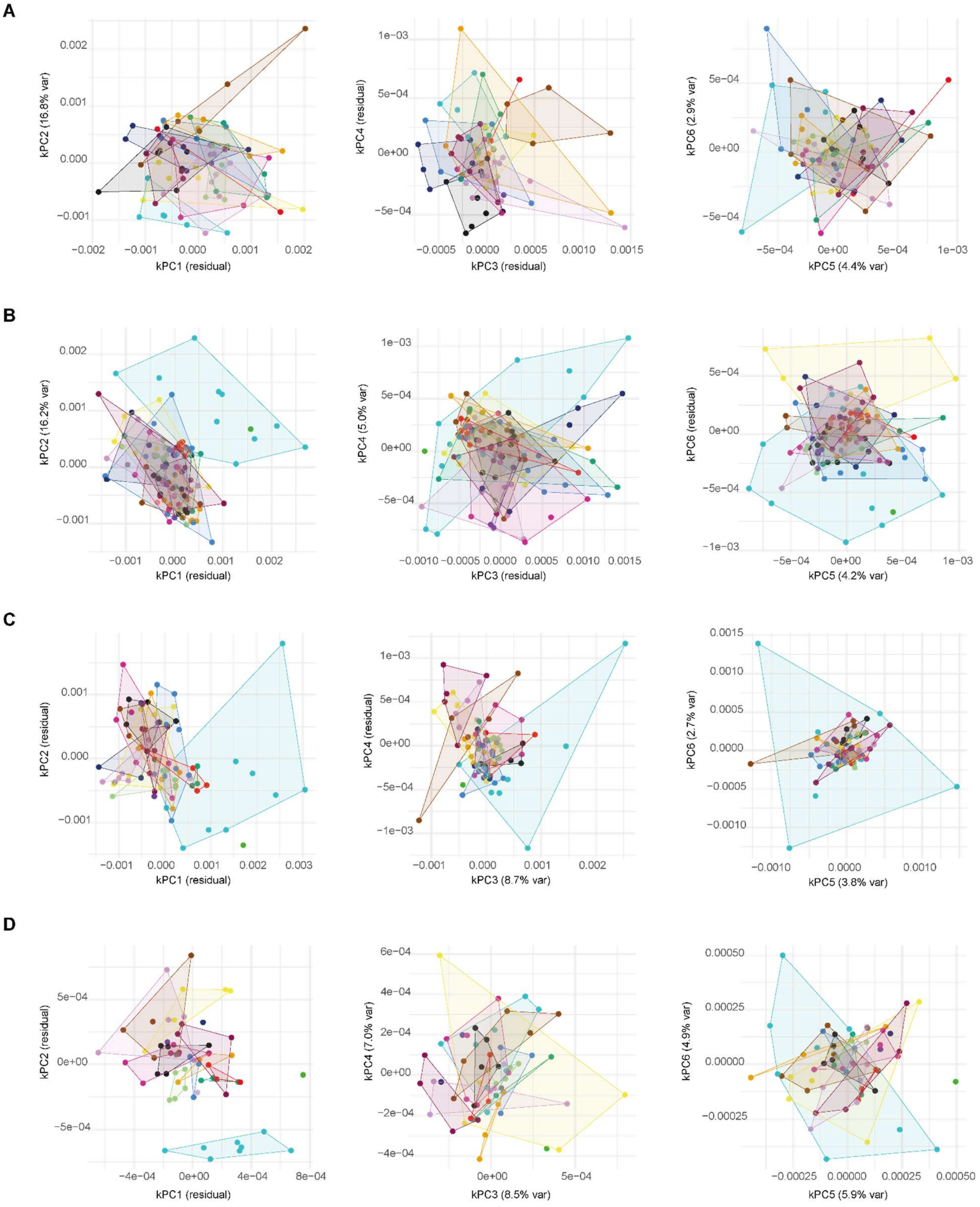

Supplementary figure 6: Partial least squares analyses for whole datasets **A** Scapula vs humerus **B** Scapula vs radius **C** Scapula vs ulna **D** Humerus vs radius **E** Humerus vs ulna **F** Radius vs ulna **G** Pelvic girdle vs femur **H** Pelvic girdle vs tibia **I** Pelvic girdle vs Fibula **J** Femur vs tibia **K** Femur vs fibula **L** Tibia vs fibula

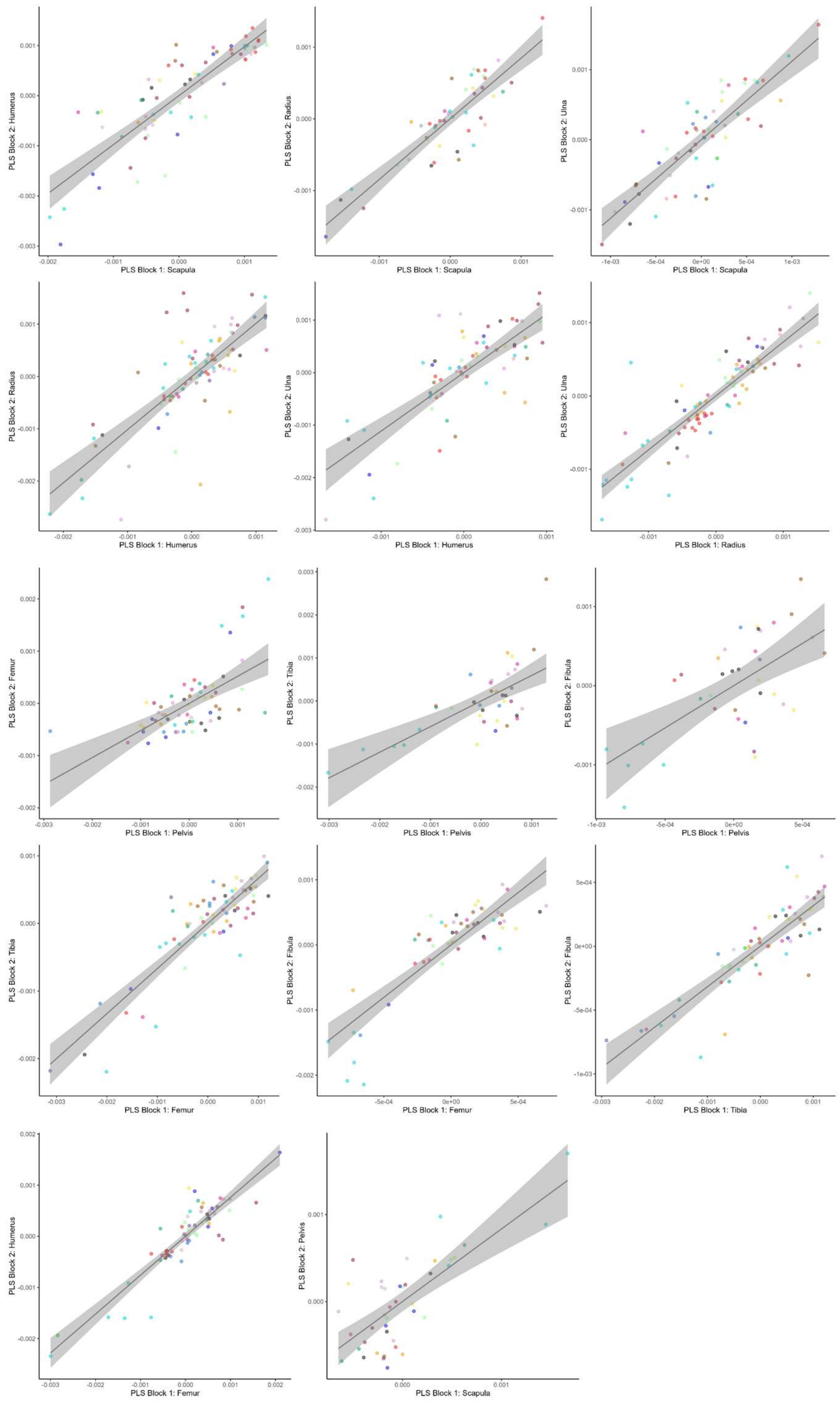

Supplementary figure 7: Instances of limb pathologies (carpal valgus, osteosarcoma, coronoid disease, luxating patella, hip dysplasia, elbow dysplasia, limb deformity & indeterminate lameness) recorded for each individual at time of scan projected onto kPC1-kPC2 plots. **A** Scapula **B** Humerus **C** Radius **D** Ulna **E** Pelvic girdle **F** Femur **G** Tibia **H** Fibula

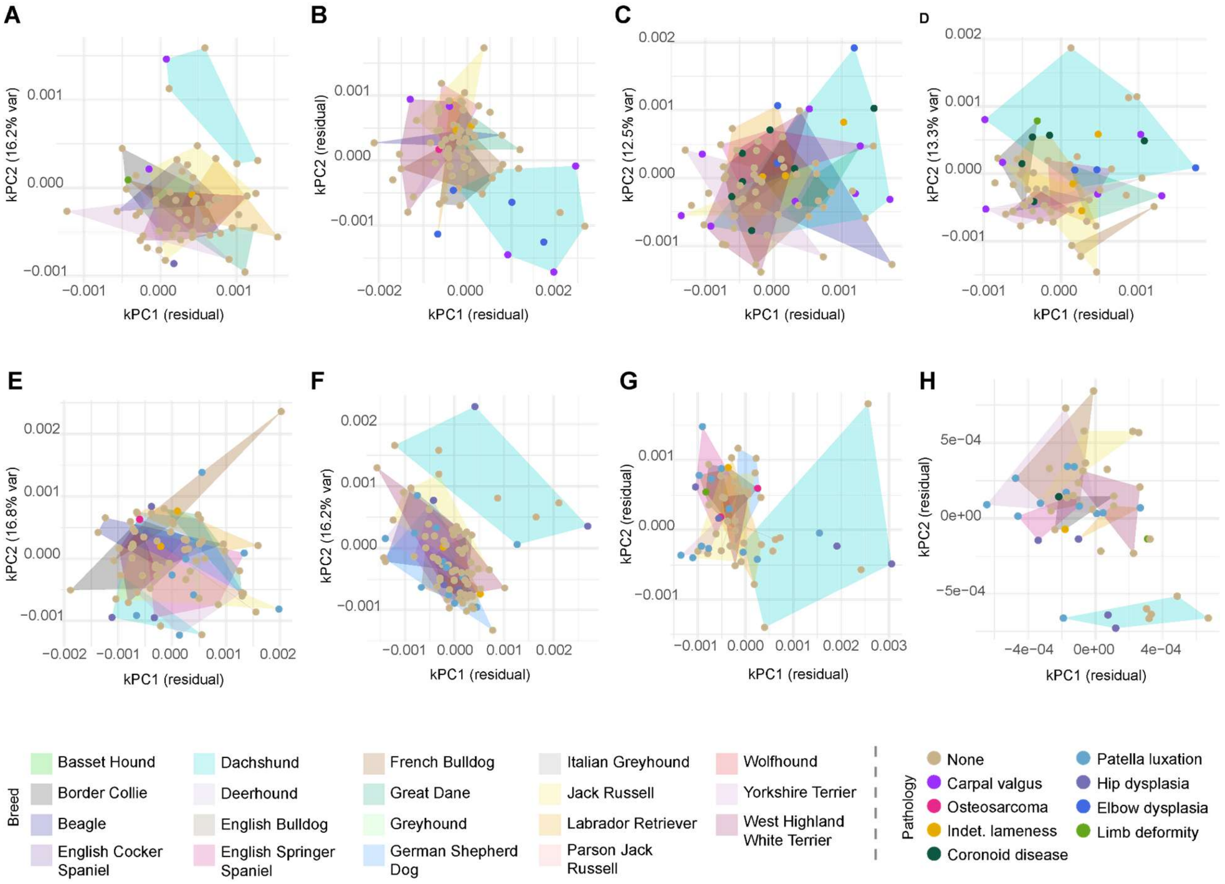

Supplementary figure 8: Body mass of individual at time of scan projected onto kPC1-kPC2 plots. **A** Scapula **B** Humerus **C** Radius **D** Ulna **E** Pelvic girdle **F** Femur **G** Tibia **H** Fibula

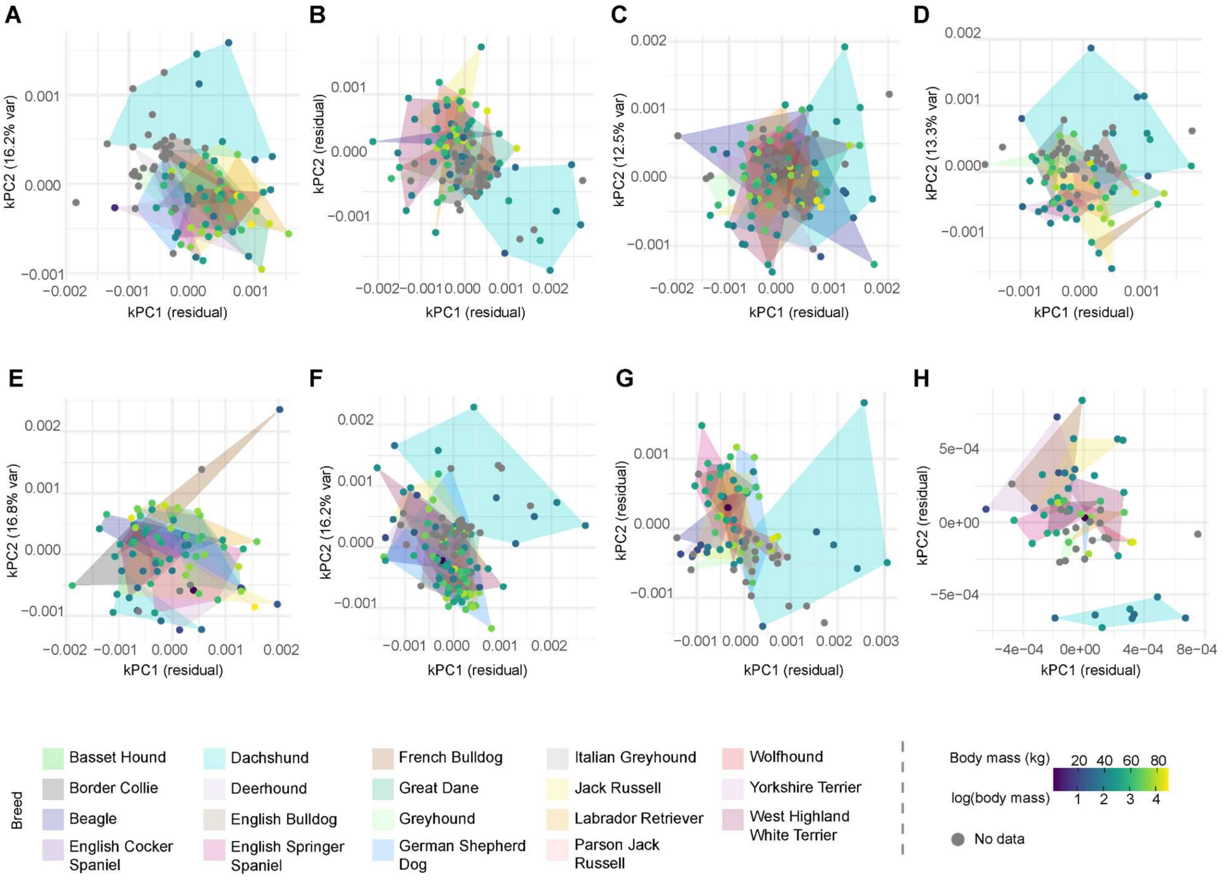

Supplementary figure 9: Age of individual at time of scan projected onto kPC1-kPC2 plots. **A** Scapula **B** Humerus **C** Radius **D** Ulna **E** Pelvic girdle **F** Femur **G** Tibia **H** Fibula

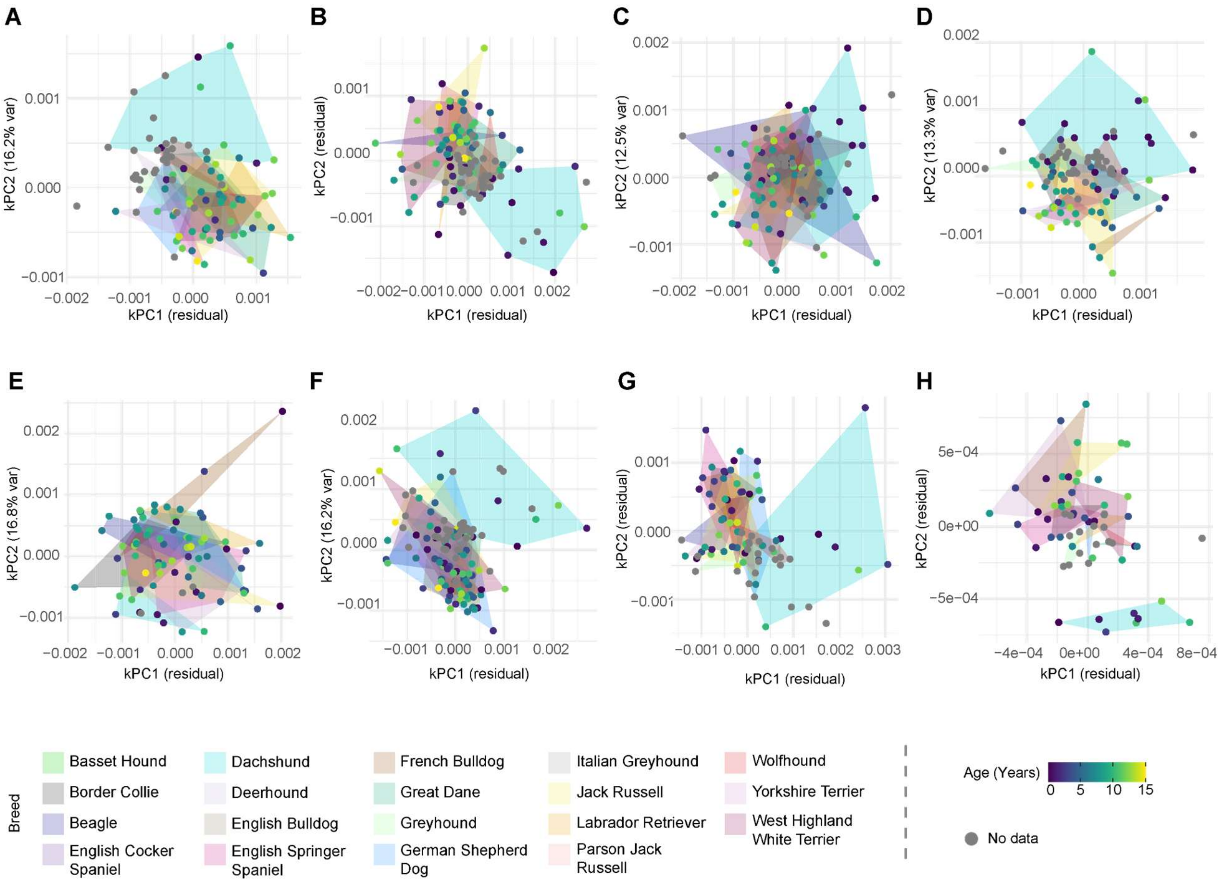

Supplementary figure 10: Change in dachshund femur morphology over time, with modern Beagle (n=1) shown for reference.

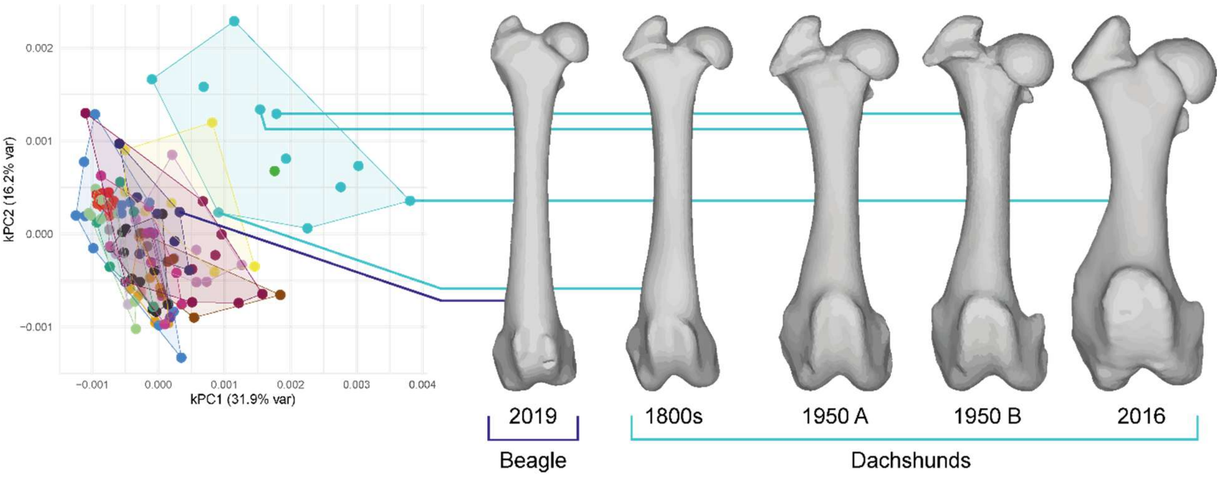

Table S1: Specimen information

Provided as excel file

Table S2: Description of landmarks used to align elements. Landmark number corresponds to Supplementary Figure 1.

| Element | Landmark | Description |
|---|---|---|
| **Scapula** | 1 | Posterior aspect of glenoid fossa border |
| **Scapula** | 2 | Ventro-medial tip of coracoid process |
| **Scapula** | 3 | Ventral tip of acromion |
| **Scapula** | 4 | Point of dorsal border at which the long axis of the scapular spine intersects |
| **Scapula** | 5 | Dorso-lateral tip of caudal angle |
| **Humerus** | 1 | Anterior point of the head |
| **Humerus** | 2 | Posterior-most point on the lateral condyle |
| **Humerus** | 3 | Posterior-most point on the medial condyle |
| **Humerus** | 4 | Proximal-most point of the olecranon fossa |
| **Humerus** | 5 | Inferior extent of the head |
| **Humerus** | 6 | Inferior-most point on the medial condyle |
| **Humerus** | 7 | Inferior-most point on the lateral condyle |
| **Radius** | 1 | Anterior-most point on head |
| **Radius** | 2 | Medial point of radial tuberosity |
| **Radius** | 3 | Proximal extent of articular facet for ulna |
| **Radius** | 4 | Distal extent of articular facet for ulna |
| **Radius** | 5 | Distal tip of the styloid process |
| **Ulna** | 1 | Anterior tip of anconeal process |
| **Ulna** | 2 | Lateral coronoid process |
| **Ulna** | 3 | Medial coronoid process |
| **Ulna** | 4 | Distal tip of styloid process |
| **Ulna** | 5 | Posterio-lateral extent of distal articular facet for radius |
| **Pelvic girdle** | 1 | Ventral extent of right acetabular margin |
| **Pelvic girdle** | 2 | Ventral extent of left acetabular margin |
| **Pelvic girdle** | 3 | Lateral extent of left ischiatic tuberosity |
| **Pelvic girdle** | 4 | Lateral extent of right ischiatic tuberosity |
| **Pelvic girdle** | 5 | Right iliopublic eminence |
| **Femur** | 1 | Center of the diaphysis in posterior view after the less trochanter |
| **Femur** | 2 | Posterior-most point on the lateral epicondyle |
| **Femur** | 3 | Posterior-most point on the medial epicondyle |
| **Femur** | 4 | Inferior-most point on the intercondylar fossa |
| **Femur** | 5 | Caudal extent of the pectineal line |
| **Femur** | 6 | Inferior-most point on the medial epicondyle |
| **Femur** | 7 | Inferior-most point on the lateral epicondyle |
| **Tibia** | 1 | Distal tip of tibial tuberosity |
| **Tibia** | 2 | Proximal tip of intercondylar eminence |
| **Tibia** | 3 | Proximal tip of lateral condyle |
| **Tibia** | 4 | Distal tip of medial malleolus |
| **Tibia** | 5 | Proximal tip of fibula notch |
| **Fibula** | 1 | Distal tip of lateral malleolus |
| **Fibula** | 2 | Cranial tip of fibula head |
| **Fibula** | 3 | Caudal extent of boundary of facet for talus and tibial notch |
| **Fibula** | 4 | Lateral tip of lateral malleolus |
| **Fibula** | 5 | Tip of apex |

Table S3: Parameters used in each Deformetrica run

| Element | Control points | Kernel width (mm) |
|---|---|---|
| Scapula | 512 | 1.33 |
| Humerus | 504 | 1 |
| Radius | 580 | 0.682 |
| Ulna | 520 | 0.78 |
| Pelvis | 504 | 1.75 |
| Femur | 525 | 0.85 |
| Tibia | 550 | 0.81 |
| Fibula | 552 | 0.426 |

Table S4: Ellipsoid volume (original and scaled). BC = Border collie, CSp = Cocker spaniel, DH = Scottish deerhound, EB = English bulldog, ESS = English springer spaniel, FB = French bulldog, GD = great dane, GH = greyhound, GSD = German shepherd dog, ItalianGH = Italian greyhound, JR = Jack Russell, Lab = Labrador retriever, PJR = Parson Jack Russell, WH = Irish Wolfhound, WHWT = West Highland white terrier.

**Original ellipsoid volumes**

| | Scapula | Humerus | Radius | Ulna | Pelvis | Femur | Tibia | Fibula |
|---|---|---|---|---|---|---|---|---|
| **BC** | 0.17 | 0.21 | 0.28 | 0.09 | 0.35 | 0.21 | 0.11 | 0.01 |
| **BassetHound** | NA | NA | NA | NA | NA | NA | NA | NA |
| **Beagle** | 0.18 | 0.21 | 3.45 | 0.03 | 0.4 | 1.65 | NA | NA |
| **CSp** | NA | NA | NA | NA | NA | NA | NA | NA |
| **DH** | 0.07 | NA | NA | NA | NA | NA | NA | NA |
| **Dachshund** | 0.35 | 2.15 | 3.26 | 1.7 | 0.24 | 3.45 | 5.95 | 0.02 |
| **EB** | NA | NA | NA | NA | NA | NA | NA | NA |
| **ESS** | 0.16 | 0.1 | 0.67 | 0.3 | 0.56 | 0.49 | 0.37 | 0.02 |
| **FB** | NA | 0.73 | 1.39 | NA | 0.44 | 0.24 | 0.5 | 0.04 |
| **GD** | 0.54 | 0.84 | 0.88 | 0.05 | 0.19 | 2.3 | NA | NA |
| **GH** | 0.2 | 0.01 | 0.06 | 0.05 | 0.07 | 0.13 | 0.03 | 0 |
| **GSD** | 0.34 | 0.02 | 0.03 | 0.02 | 1.03 | 1.93 | 0.29 | NA |
| **ItalianGH** | NA | NA | NA | NA | NA | NA | NA | NA |
| **JR** | 0.26 | 0.05 | 0.91 | NA | 0.15 | 1.69 | 0.64 | 0.05 |
| **Lab** | 0.24 | 0.4 | 0.51 | 0.2 | 0.42 | 0.16 | 0.14 | 0 |
| **PJR** | NA | NA | NA | NA | NA | NA | NA | NA |
| **WH** | 0.07 | 0.02 | 0.06 | 0.03 | NA | 0.02 | 0.23 | NA |
| **WHWT** | 0.41 | 0.89 | 0.69 | 0.03 | 0.21 | 0.68 | 0.22 | 0.01 |
| **Yorkie** | 0.22 | 0.32 | 0.41 | 0.27 | 1.51 | 0.42 | 0.07 | 0.14 |

**Scaled ellipsoid volumes**

| | Scapula | Humerus | Radius | Ulna | Pelvis | Femur | Tibia | Fibula |
|---|---|---|---|---|---|---|---|---|
| **BC** | -0.57 | -0.41 | -0.61 | -0.33 | -0.28 | -0.76 | -0.39 | -0.51 |
| **BassetHound** | NA | NA | NA | NA | NA | NA | NA | NA |
| **Beagle** | -0.5 | -0.41 | 2.2 | -0.45 | -0.15 | 0.58 | NA | NA |
| **CSp** | NA | NA | NA | NA | NA | NA | NA | NA |
| **DH** | -1.32 | NA | NA | NA | NA | NA | NA | NA |
| **Dachshund** | 0.77 | 2.83 | 2.03 | 2.95 | -0.54 | 2.26 | 3 | -0.28 |
| **EB** | NA | NA | NA | NA | NA | NA | NA | NA |
| **ESS** | -0.65 | -0.6 | -0.27 | 0.1 | 0.23 | -0.5 | -0.24 | -0.28 |
| **FB** | NA | 0.46 | 0.37 | NA | -0.06 | -0.74 | -0.16 | 0.18 |
| **GD** | 2.19 | 0.64 | -0.08 | -0.41 | -0.66 | 1.19 | NA | NA |
| **GH** | -0.35 | -0.75 | -0.81 | -0.41 | -0.95 | -0.84 | -0.43 | -0.74 |
| **GSD** | 0.69 | -0.73 | -0.83 | -0.47 | 1.37 | 0.84 | -0.28 | NA |
| **ItalianGH** | NA | NA | NA | NA | NA | NA | NA | NA |
| **JR** | 0.1 | -0.68 | -0.05 | NA | -0.76 | 0.62 | -0.08 | 0.41 |
| **Lab** | -0.05 | -0.1 | -0.41 | -0.11 | -0.11 | -0.81 | -0.37 | -0.74 |
| **PJR** | NA | NA | NA | NA | NA | NA | NA | NA |
| **WH** | -1.32 | -0.73 | -0.81 | -0.45 | NA | -0.94 | -0.32 | NA |
| **WHWT** | 1.22 | 0.72 | -0.25 | -0.45 | -0.61 | -0.33 | -0.32 | -0.51 |
| **Yorkie** | -0.2 | -0.23 | -0.5 | 0.04 | 2.53 | -0.57 | -0.41 | 2.46 |

Table S5: Statistics from regression of kPCs against centroid size. Significant ($P < 0.05$) relationships only.

| Element | kPC | pval | adj-rsq |
|---|---|---|---|
| Scapula | kPC1 | <0.001 | 0.36 |
| Scapula | kPC2 | 0.001 | 0.11 |
| Scapula | kPC3 | 0.008 | 0.07 |
| Scapula | kPC4 | 0.008 | 0.07 |
| Scapula | kPC5 | 0.003 | 0.09 |
| Humerus | kPC1 | <0.001 | 0.42 |
| Humerus | kPC2 | <0.001 | 0.14 |
| Humerus | kPC3 | <0.001 | 0.11 |
| Radius | kPC1 | <0.001 | 0.54 |
| Radius | kPC5 | 0.006 | 0.06 |
| Ulna | kPC1 | <0.001 | 0.62 |
| Ulna | kPC4 | 0.035 | 0.04 |
| Ulna | kPC10 | 0.045 | 0.03 |
| Pelvic girdle | kPC1 | <0.001 | 0.32 |
| Pelvic girdle | kPC3 | <0.001 | 0.15 |
| Pelvic girdle | kPC4 | 0.010 | 0.07 |
| Femur | kPC1 | <0.001 | 0.50 |
| Femur | kPC3 | <0.001 | 0.10 |
| Femur | kPC6 | 0.006 | 0.05 |
| Tibia | kPC1 | <0.001 | 0.49 |
| Tibia | kPC2 | 0.051 | 0.04 |
| Tibia | kPC4 | 0.003 | 0.10 |
| Fibula | kPC1 | <0.001 | 0.62 |
| Fibula | kPC2 | 0.048 | 0.05 |

Table S6: Integration statistics

| | Scapula | Humerus | Radius | Ulna | Pelvis | Femur | Tibia | Fibula |
|---|---|---|---|---|---|---|---|---|
| **Scapula** | | 0.752 | 0.869 | 0.827 | 0.747 | 0.783 | 0.884 | 0.868 |
| **Humerus** | 4.164 | | 0.775 | 0.775 | 0.694 | 0.900 | 0.963 | 0.913 |
| **Radius** | 4.871 | 4.932 | | 0.877 | 0.778 | 0.870 | 0.925 | 0.935 |
| **Ulna** | 4.884 | 4.789 | 6.548 | | 0.859 | 0.908 | 0.917 | 0.972 |
| **Pelvis** | 3.214 | 1.834 | | | | 0.617 | 0.685 | 0.639 |
| **Femur** | 4.280 | 4.891 | 4.043 | 3.699 | 2.639 | | 0.874 | 0.854 |
| **Tibia** | | | | | 2.281 | 4.899 | | 0.859 |
| **Fibula** | | | | | | 4.449 | 5.298 | |

Table S7: Results of PERMANOVA tests assessing breed, breed group, nested breed, age, body mass and recorded pathology.

Provided as excel file

Table S8: Statistics for significant ($p < 0.05$) pairwise PERMANOVAs testing for shape differences between breeds. FB = French bulldog, WH = wolfhound, ESS = English springer spaniel, GH = greyhound, JR = Jack Russell terrier, CSp = Cocker spaniel, GSD = German shepherd dog, DH = deerhound, BC = Border collie, WHWT = West Highland white terrier, GD = Great Dane, Lab = Labrador retriever.

Provided as excel file

Table S9: Results of pairwise PERMANOVAs assessing differences in the radius and humerus shape between dogs grouped by recorded pathology. CV = carpal valgus, E = elbow dysplasia, N = ‘none’, OA = osteoarthritis, L = indeterminate lameness, CD = coronoid disease.

Provided as excel file

Table S10: Results of PERMANOVAs assessing differences in the element shape between within breeds grouped by recorded pathology.

Provided as excel file
